# Enhancer RNA-based modeling of adverse events and objective responses of immunotherapy

**DOI:** 10.1101/2021.12.16.473069

**Authors:** Mengbiao Guo, Zhiya Lu, Yuanyan Xiong

**Author notes:** These authors contributed equally to this work (Mengbiao Guo, Zhiya Lu).

## Abstract

Immune checkpoint inhibitors (ICI) targeting PD-1/PD-L1 or CTLA-4 are emerging and effective immunotherapy strategies. However, ICI treated patients present heterogeneous responses and adverse events, thus demanding effective ways to assess benefit over risk before treatment. Here, by integrating pan-cancer clinical and molecular data, we tried to predict immune-related adverse events (irAEs, risk) and objective response rates (ORRs, benefit) based on enhancer RNAs (eRNAs) expression among patients receiving anti-PD-1/PD-L1 therapy. We built two effective regression models, explaining 71% variance (R=0.84) of irAEs with three eRNAs and 79% (R=0.89) of ORRs with five eRNAs. Interestingly, target genes of irAE-related enhancers, including upstream regulators of MYC, were involved in metabolism, inflammation, and immune activation, while ORR-related enhancers target *PAK2* and *DLG1* which directly participate in T cell activation. Our study provides references for the identification of immunotherapy-related biomarkers and potential therapeutic targets during immunotherapy.

## Introduction

Immune checkpoints (ICs) generally refer to key inhibitory factors of the immune system, including programmed cell death 1 (PD-1 or CD279) and its ligand programmed cell death 1 ligand 1 (PD-L1 or CD274) that control the T cell response and fate during tumor immunity [1]. In tumor samples, PD-1 and PD-L1 mainly expressed in T cells and tumor cells, respectively, and tumors exploit their interaction to escape the immune system by counteracting the stimulatory signals from the interaction between T cell receptor (TCR) and major histocompatibility complex (MHC) and other costimulatory signals [2–4].

PD-1/PD-L1 has been translated to the clinical practice, and ICI treatment targeting PD-1/PD-L1 proved to offer significant clinical benefits in many cancers, with an ORR from 20% to 50% in multiple clinical trials and for various types of cancer [5]. However, only a small subset of patients showed long-lasting remission, despite remarkable benefits of ICI therapies. Patients of some cancers were completely refractory to checkpoint blockade, occasionally leading to considerable side effects. To predict treatment benefit, PD-L1 expression was proposed as the first biomarker of anti–PD-1/PD-L1 therapy effectiveness [6], followed by tumor mutational burden (TMB) [7]. Later, microsatellite instability (MSI) [8], CD8+ T-cell abundance [9, 10], cytolytic activity [11], and intestinal microbial composition[12] were proposed to prioritize patients with potentially more treatment gains.

On the other hand, irAEs result from excessive immunity against normal organs. Most studies show that the incidence of irAEs caused by anti-PD-1/PD-L1 treatment is about 60% [13, 14]. Although nearly all organs can be affected, irAEs mostly involved the gastrointestinal tract, endocrine glands, skin, and liver [15]. In some cases, irAE can be lethal. For example, pneumonitis is the most common fatal irAE with a 10% death rate, accounting for 35% of anti-PD-1/PD-L1-related fatalities [16]. The mortality of myocarditis, the most lethal irAE, could even reach about 50% [17]. Therefore, it is important and urgent to select patients with potentially significant benefit over risk of ICI treatments based on individual molecular data. Although people have discovered several predictors of irAEs using expression of protein-coding genes [18], studying irAE-related non-coding elements would probably provide a better mechanistic understanding of why PD-1/PD-L1 pathway modulation leads to significant clinical benefit in some patients but temporary, partial, or no clinical benefit in other patients.

Recent studies found that eRNAs (non-coding RNAs) were usually transcribed from active enhancers and eRNA levels portended enhancer activities across tissues [19]. Numerous cancer-associated eRNAs have been identified and eRNAs were proposed as potential therapeutic targets [20]. Here, we comprehensively investigate the adverse events and the response rates in patients receiving anti-PD-1/PD-L1 therapies across cancer types. By integrating clinical data and molecular data, we identify predictors based on three eRNAs for predicting irAE and five eRNAs for ORR. Further exploring enhancer-target interaction identified functional genes that may help explain the overall risk or benefit of anti-PD-1/PD-L1 therapy, including MLXIPL, RAF1, MPL, PAK2, DLG1. In summary, our study reveals potential mechanisms underlying ICI therapy based on enhancer activity.

## Results

### Three eRNAs effectively predict irAE of immunotherapy

To identify factors to predict irAEs, we first examined correlations between 7 045 eRNAs and irAE RORs across 25 cancer types and found 178 eRNAs positively correlated with irAEs with nominal significance (*P*<0.05). Among these eRNAs, ENSR00000041252 showed the highest correlation (correlation R=0.68, *P*=1.6e-4; **Fig. S1A**), stronger than immune factors, including naive B cells, CD8+ T cells, macrophages M1, and T cell receptor diversity [18].

Then, we selected the top ten eRNAs (**Table S1**) to build prediction models. Multicollinearity analysis resulted in six roughly independent eRNAs, ENSR00000041252, ENSR00000326714, ENSR00000148786, X14.65054944.65060944, ENSR00000118775, and ENSR00000242410 (**Fig. 1A** and **Fig. 1B**). Next, we obtained 15 significant bivariate regression models using the irAE-correlated enhancers. Correlation between the observed and predicted irAE ROR values showed that the combination ENSR00000148786 + ENSR00000005553 achieved the best predictive performance (R=0.79, *P*=3.1e-6; **Fig. S1B**). Further increasing model factors resulted in the optimal tri-variate model, ENSR00000041252 + ENSR00000148786 + ENSR00000005553, with the strongest correlation (R=0.84, *P*=2.1e-6; **Fig. 1C**). Of note, no improvement was observed after adding the two protein-coding genes (LCP1 and ADPGK) from a model reported previously [18], suggesting the independence of our model. Although showing slightly lower performance than the previous protein-coding gene model (LCP1+ADPGK), our enhancer-based model, explaining 71% (R-squared, R=0.84) of irAE variance, demonstrated that eRNAs alone can effectively predict irAEs.

**Fig.1.**
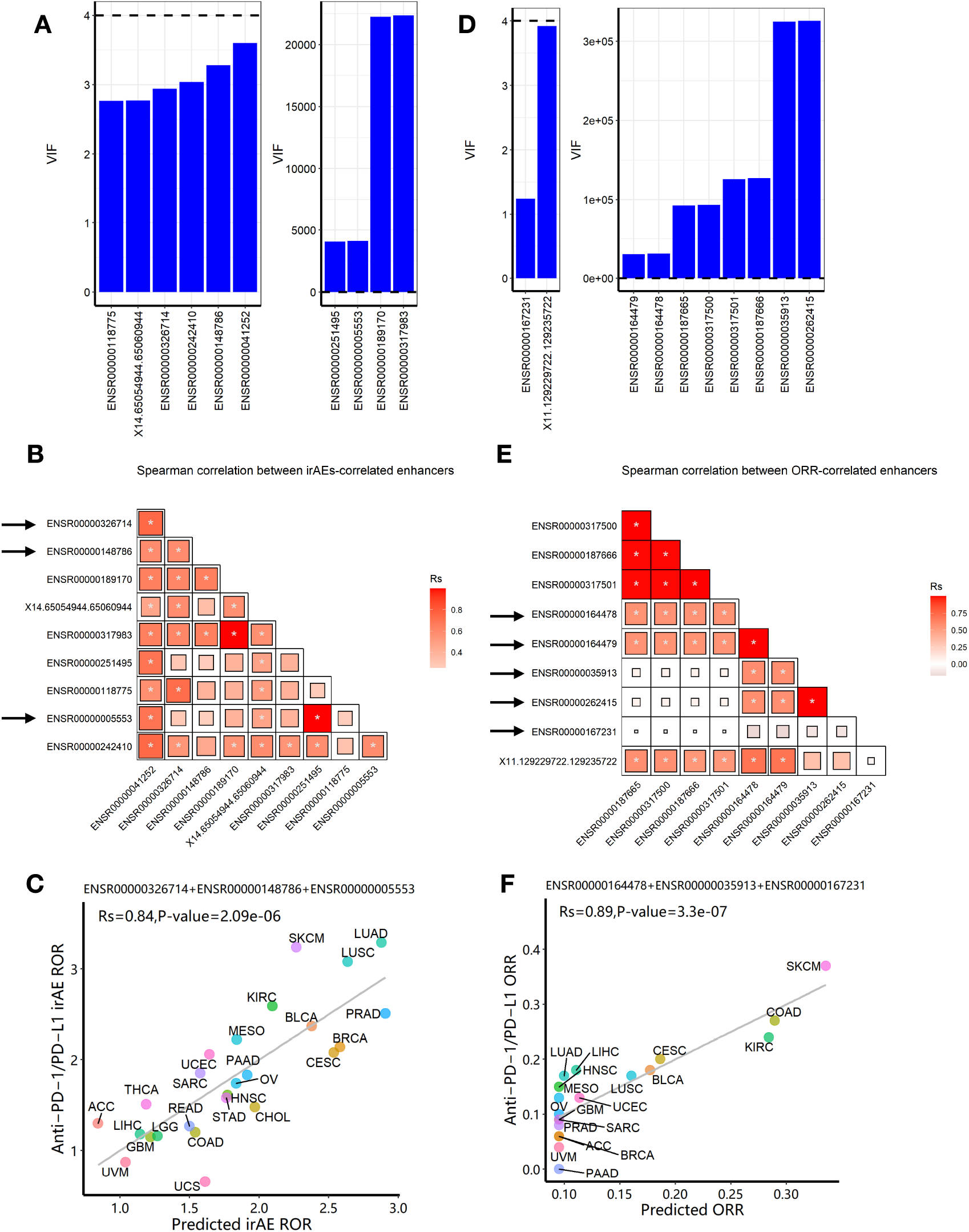
Construction of eRNA-based prediction models for irAE ROR (risk) and ORR (benefit) of immunotherapy. (**A**) Multicollinearity (VIF) analysis for top ten eRNA expression in predicting irAEs. Six eRNAs showed no multicollinearity, while 4 eRNAs showed strong multicollinearity. (**B**) Spearman correlation between irAE-correlated eRNAs. Pairwise Spearman correlation (Rs) of expression level between candidate eRNAs. The shade of the square indicates the Rs, and the size indicates P-value (* indicates statistical significance *P*<0.05). (**C**) Combined effect of ENSR00000326714, ENSR00000148786 and ENSR00-000005553 trivariate model of predicting irAEs (R=0.84, *P*=2.1e-6). The equation of the best trivariate model is 0.1912*ENSR00000005553+0.4097*ENSR000-00326714+0.1953*ENSR00000148786+0.2942. (**D**) Multicollinearity analysis for top ten eRNA expression in predicting ORR. Two eRNAs showed no multicollinearity, while 8 eRNAs showed strong multicollinearity. (**E**) Spearman correlation between ORR-correlated eRNAs. Spearman correlation (*Rs*) of expression level was calculated between two candidate eRNAs. The shade of the square indicates the *Rs*, and the size indicates P-value (* indicates statistical significance *P*< 0.05). (**F**) Combined effect of ENSR00000164478, ENSR00000035913 and ENSR000-00167231 trivariate model of predicting ORR (R=0.89, *P*=3.3e-7). The equation of the best trivariate model is 0.0953+0.0649* ENSR00000164478+0.0032* ENSR00000035913+0.1687* ENSR00000167231. irAE, immune-related adverse events; ROR, reporting odds ratio; ORR, objective response rates; LUAD, lung adenocarcinoma; SKCM, skin cutaneous melanoma; LUSC, lung squamous cell carcinoma; KIRC, kidney renal clear cell carcinoma; PRAD, prostate adenocarcinoma; BLCA, bladder urothelial carcinoma; MESO, mesothelioma; BRCA, breast invasive carcinoma; CESC, cervical squamous cell carcinoma and endocervical adenocarcinoma; UCEC, uterine corpus endometrial carcinoma; SARC, sarcoma; ESCA, esophageal carcinoma; PAAD, pancreatic adenocarcinoma; OV, ovarian serous cystadenocarcinoma; HNSC, head and neck squamous cell carcinoma; STAD, stomach adenocarcinoma; THCA, thyroid carcinoma; CHOL, cholangiocarcinoma; ACC, adrenocortical carcinoma; READ, rectum adenocarcinoma; COAD, colon adenocarcinoma; LIHC, liver hepatocellular carcinoma; LGG, brain lower-grade glioma; GBM, glioblastoma multiforme; UVM, uveal melanoma; UCS, uterine carcinosarcoma.

### Five eRNAs effectively predict immunotherapy benefit

Similarly, to identify factors to predict ORRs, we identified 28 out of 7 045 eRNAs positively correlated with ORR (*P*<0.05; the best one ENSR00000187665 shown in **Fig. S1C**). Based on the top ten eRNAs (**Table S2**), after multicollinearity analysis (**Fig. 1D** and **Fig. 1E**), two bivariate models achieved better predictive performance than single-eRNA models (one shown in **Fig. S1D**; R=0.82, *P*=2.0e-5). Further adding model factors resulted in four equally-efficient optimal trivariate models (involving five key eRNAs, **Table S3**) for ORR prediction were able to effectively predict the efficacy of anti–PD-1/PD-L1 treatments. One example, ENSR00000164478 + ENSR00000035913+ ENSR00000167231, was shown in **Fig. 1F** (R=0.89, *P*=3.3e-7).

### Enhancer-target networks of irAE and ORR-associated enhancers

Enhancers were assumed to affect irAEs or ORRs by activating target genes through long-range interactions. We downloaded enhancer-target interaction data[21] and obtained putative targets of our enhancers. Two eRNAs (ENSR00000262415 and ENSRO0000167231) were excluded from downstream analysis due to lack of any annotated target gene. eRNA-target networks showed that these enhancers independently regulated a specific groups of targets (**Fig. 2A** and **Fig. 2B**, note that ENSR00000164478 and ENSR00000164479 located to the same genomic region), indicating that each irAE-related enhancer was involved in different regulatory modules. Similarly, protein-protein interaction (PPI) analysis revealed that an independent network was controlled by each enhancer (**Fig. 1C** and **Fig. 1D**). In these PPI networks, genes located in the center (such as BCL7B, TBL2, and NAP1L4) might be vital regulators of irAEs or ORRs.

**Fig.2.**
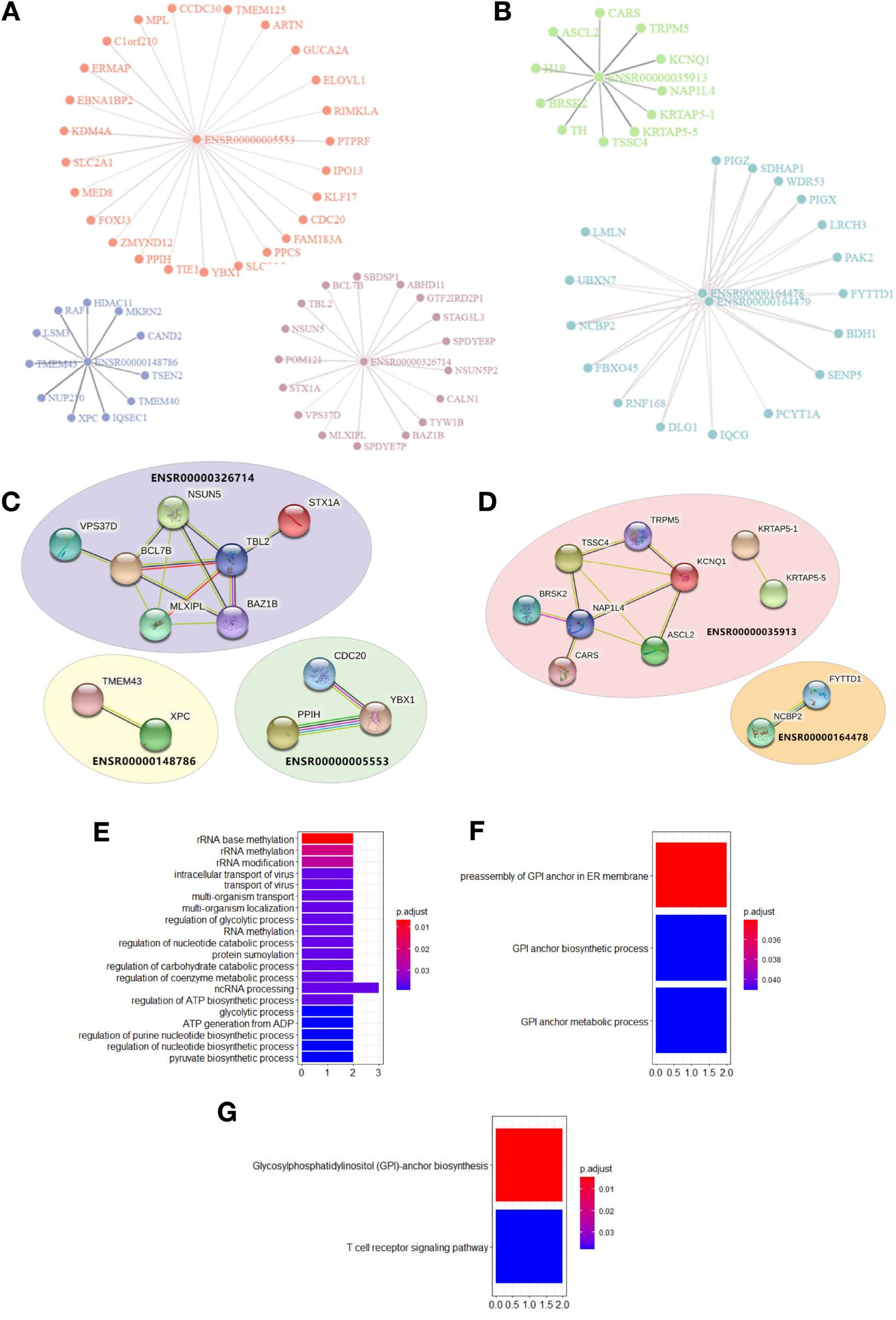
Visualization of enhancer-target interaction network and functional enrichment. **(A)** target genes of irAE-related enhancers ENSR00000005553, ENSR00000326714, and ENSR00000148786. **(B)** target genes of ORR-related enhancers ENSR00000164478, ENSR00000164479, and ENSR00000035913. (**C**) Protein-Protein Interaction (PPI) network for target genes of irAE-related enhancer ENSR00000326714, ENSR00000148786, ENSR00000005553; and their corresponding PPI of targets in irAE ROR model. **(D)** PPI network for targets of ORR-related enhancers ENSR00000035913, ENSR000-00164478. (**E**) GO enrichment of genes regulated by irAE-correlated enhancer ENSR00000326714. (**F**) GO enrichment of genes regulated by ORR-correlated enhancer ENSR00000164478. (**G**) KEGG pathway enrichment of genes regulated by ORR-correlated enhancer ENSR00000164478.

### Enhancer targets reveal metabolic and inflammatory genes involved in irAEs

Next, we downloaded gene sets from COSMIC[22] and oncoKB[23] and examined our eRNA targets in known oncogenic signaling pathways using cBioPortal[24, 25]. We found that some eRNA targets were known cancer genes relevant to tumor immunity, including *MLXIPL, MPL, RAF1*, and *XPC. RAF1* was annotated as an oncogene and participated in the RTK-RAS signaling pathway (**Fig. S2A**) and MLXIPL was involved in MYC signaling pathway (**Fig. S2B**). A previous work[26] shows RAF1 can activate MAPK1 and NF-κB pathways to regulate genes involved in inflammation. Therefore, RAF1 may enhance immunoreaction and subsequently cause irAEs via Natural Killer cell-mediated cytotoxicity, T cell receptor signaling pathway, and B cell receptor signaling pathway.

Interestingly, we found that ENSR00000326714 targets were enriched in a large number of metabolic and biosynthesis processes (**Fig. 2E**). This was reminiscent of some types of adverse events, such as diabetes[16], due to metabolic disturbances or metabolic disorders. Specifically, the core network of ENSR00000326714 targets consists of seven metabolic and inflammatory genes, namely, BAZ1B, BCL7B, TBL2, MLXIPL, NSUN, STX1A, and VPS37D. Among them, BAZ1B, BCL7B, TBL2 and MLXIPL are pleiotropic genes for lipids and inflammatory markers in the liver[27]. Of note, MLXIPL encodes the carbohydrate-responsive element-binding protein (ChREBP), which mediates glucose homeostasis and liver lipid metabolism. ChREBP was also associated with up-regulation of several cytokines (TNF-α, IL-1β, and IL-6) in patients with type 2 diabetes mellitus, promoting the inflammatory responses and apoptosis of mesangial cells[28]. STX1A encodes a member of the syntaxin superfamily, syntaxin 1A. It contributes to neural function in the central nervous system by regulating transmitter release[29]. As a kind of target-SNAP receptor (t-SNAREs), it is involved in insulin exocytosis[30]. Severely reduced islet syntaxin 1A level was reported to contribute to insulin secretory deficiency[31]. Given that diabetes and hepatitis account for ~30% of immune-related adverse events[16], we speculate that ENSR00000326714 augmented the expression of the these genes, subsequently triggering inflammation and other toxic effects on these patients.

### ORR enhancers reveal immune activation genes for immunotherapy benefit

We also analyzed target genes of ORR-predictable eRNAs (**Fig. 2B**), which included three types of genes. PAK2, LMLN, DLG1, ASCL2, SENP5, IQCG, and BRSK2 are related to cell cycle, cell division, and differentiation. PIGZ, PIGX, PCYT1A, CARS, and BDH1 are metabolic genes; TRPM5, KCNQ1, and FYTTD1 are responsible for cellular transport and signal transduction. In particular, target genes of ORR-related ENSR00000164478 were enriched in glycosylphosphatidylinositol (GPI)-anchor biosynthesis (FDR=4.73 × 10^−3^) (**Fig. 2F**) and T-cell receptor signaling (FDR=3.78×10^−2^), among other enriched pathways (**Fig. 2G**).

Furthermore, PAK2 and DLG1 directly took part in the T cell activation pathway, which explains their connection with ORR. P21 (RAC1) activated kinase 2 (PAK2) has been reported as a key signaling molecule in the differentiation of T cells. PAK2 is essential in T cell development and differentiation[32], indicating its potential function in T cell-initiated autoimmunity. DLG1 encodes a multi-domain scaffolding protein from the membrane-associated guanylate kinase family, which has been shown to regulate the antigen receptor signaling and cell polarity in lymphocytes, involved in activation and proliferation of T cells[33, 34]. Our results provide more support for the T cells as the regulators in immune responses during immune checkpoint blockade therapy.

Lastly, PIGZ encodes a protein that is previously identified as an immune-associated prognosis signature[35]. However, knowledge of the relationship between PIGZ and the immune system is still poorly established. The association between PIGZ expression and immune benefits during anti-PD1/PDL1 immunotherapy needs further elucidation.

## Discussions

In this work, we presented a preliminary evaluation of the different enhancer-target interactions associated with anti–PD-1/PD-L1 immunotherapy across tumor types, and successfully identify potential enhancer-based biomarkers of risk and beneficial response. We suggest that, during immunotherapy, enhanced expression of inflammatory factors including MLXIPL, STX1A, and RAF1 may lead to a higher risk of irAEs, while strengthening immune activation factors including PAK2 and DLG1 may improve anti-tumor immunity. Besides, we discovered many other cancer-related, metabolic, signaling or regulatory genes possess predictive potential, which warrants further investigation.

Several limitations remain for future work and our results need to be carefully interpreted. First, the majority of data are collected from previous individual studies[21], introducing inherent limitations of their work. Second, there are inevitable flaws of modeling as well, due to the low expression level of eRNA and small sample size. The overall quality of predictive models of ORR is inferior to those of irAEs, probably due to a smaller sample size as well as larger sparsity of ORR data. Finally, since results in this project are mainly based on computational predictions and the support of existing literature, our findings need further experimental validation. A larger dataset is required to comprehensively model side effects or immune response as well.

## Methods

### Data collection

To quantify the risk of immune-related adverse events (irAEs), reporting odds ratio (ROR) was calculated as previously described [36]. The anti-PD1/PD-L1 irAE ROR and ORR values across different cancer types were collected from previous studies [10, 18]. RNA-seq expression data (RSEM normalized counts, log2-transformed) across 25 TCGA cancers were downloaded from the UCSC Xena platform (http://xena.ucsc.edu/). Expression levels of selected genes were extracted for downstream analysis, and the average value was calculated for each TCGA cohort. We downloaded eRNA expression levels and enhancer-target associations for 7 045 enhancer RNAs in ~7,300 samples from the eRic database [21] (https://hanlab.uth.edu/eRic/). Mean eRNA expression (log2-transformed RPM values) were used. Similar to gene expression, we averaged the expression level of each eRNA for each cancer.

### Prediction model construction

First, the top ten eRNAs were selected based on correlation between eRNA and irAE or ORR. Before constructing bivariate models, the variance inflation factor[37] (VIF) of these ten eRNAs was calculated to evaluate the multicollinearity. Generally, we set the threshold of VIF value to 4 (a VIF value greater than 10 will be considered serious multicollinearity). The optimal prediction model was obtained by step-wise addition of model factors (eRNA) and evaluate the correlation between predicted and observed patient risk or benefits.

### Bioinformatics tools

We used the protein-protein interaction (PPI) database STRING[38] (v11, https://string-db.org) to investigate selected eRNA target genes. Basic GO and KEGG term enrichment and visualization were conducted with the R package clusterProfiler[39] (v3.14.3). Extensive functional annotation of eRNA target genes were performed with DAVID [40] (v6.8) (https://david.ncifcrf.gov/). To verify cancer-related function for genes of interest, a credible set of 723 cancer genes was downloaded from the Cancer Gene Census (CGC) project of the COSMIC[22] repository (https://cancer.sanger.ac.uk/cosmic/). Another database oncoKB[23] (https://oncokb.org/), which has a list of 1,064 cancer genes, was added as a supplement to COSMIC CGC genes. Oncogenic signaling pathways were provided by the cBioPortal database[24] (http://www.cbioportal.org/). Statistical analysis and visualization were performed in R (v3.6.3) using packages ggplot2 (v3.3.2), networkD3 (v0.4). For novel candidates, we used three types of biological interpretation (Gene Oncology, Pathways, and Protein-Protein Interaction) to obtain biological knowledge.

### Statistical methods

We employed an approach as described previously [10, 18] to evaluate the correlation between eRNAs and irAE RORs or ORRs. Linear-regression models for predicting irAE ROR or ORR across cancer types, was constructed by the R function lm, and the performance of the prediction was estimated based on Spearman rank correlation, using the R package psych (v2.0.12). To compare the goodness of fit between different models, a log-likelihood ratio test was performed using the R package lmtest (v0.9). We compute variance inflation factor (VIF) to assess multicollinearity using the vif function from the R package car (v3.0) to exclude combinations containing highly correlated factors.

## Acknowledgements

The research has been supported by National Natural Science Foundation of China (NSFC) (Grant 31571350, U1611265, and 31871323). The results shown here are in whole or part based upon data generated by the TCGA or CPTAC Research Network.

## Author contributions

YYX and MBG conceived and supervised the study. ZYL, YYX, and MBG performed the analysis. MBG drafted the manuscript with assistance from ZYL. YYX reviewed the manuscript. All authors approved the final manuscript.

