## Supplementary material for "Enhancer RNA-based modeling of adverse events and objective responses of immunotherapy": Supllementary Figures S1-2 and Tables S1-3

### Supplementary Information

#### Supplementary Tables

**Table S1. Top ten significant eRNAs associated with irAE.**

| eRNA | R | *P*-value |
|---|---|---|
| ENSR00000041252 | 0.68 | 1.64E-04 |
| ENSR00000326714 | 0.66 | 3.17E-04 |
| ENSR00000148786 | 0.66 | 3.75E-04 |
| ENSR00000189170 | 0.62 | 8.95E-04 |
| X14.65054944.65060944 | 0.62 | 1.01E-03 |
| ENSR00000317983 | 0.62 | 1.06E-03 |
| ENSR00000251495 | 0.61 | 1.31E-03 |
| ENSR00000118775 | 0.60 | 1.46E-03 |
| ENSR00000005553 | 0.60 | 1.59E-03 |
| ENSR00000242410 | 0.60 | 1.62E-03 |

**Table S2. Top ten significant eRNAs associated with ORR.**

| Factor name | eRNA | R | *P*-value |
|---|---|---|---|
| *x1* | ENSR00000187665 | 0.59 | 7.46E-03 |
| *x2* | ENSR00000317500 | 0.59 | 7.46E-03 |
| *x3* | ENSR00000187666 | 0.55 | 1.37E-02 |
| *x4* | ENSR00000317501 | 0.55 | 1.37E-02 |
| *x5* | ENSR00000164478 | 0.51 | 2.44E-02 |

| | | | |
|---|---|---|---|
| *x6* | ENSR00000164479 | 0.51 | 2.44E-02 |
| *x7* | ENSR00000035913 | 0.51 | 2.45E-02 |
| *x8* | ENSR00000262415 | 0.51 | 2.45E-02 |
| *x9* | ENSR00000167231 | 0.50 | 2.76E-02 |
| *x10* | X11.129229722.129235722 | 0.50 | 2.78E-02 |

**Table S3. Combined effects of the best trivariate ORR models.** *x5* ENSR00000164478, *x6* ENSR00000164479, *x7* ENSR00000035913, *x8* ENSR00000262415, *x9* ENSR-00000167231.

| Trivariate model | Rs | P-value |
|---|---|---|
| *x5+x7+x9* | 0.89 | 3.31E-07 |
| *x6+x7+x9* | 0.89 | 3.31E-07 |
| *x5+x8+x9* | 0.89 | 3.00E-07 |
| *x6+x8+x9* | 0.89 | 3.00E-07 |

#### Supplementary Figures

**Fig. S1. Correlation between eRNAs and irAE or ORR.**

(**A**) Spearman correlation between ENSR00000041252 expression and irAE ROR (R=0.68, *P*=1.6e-4). (**B**) Combined effect of ENSR00000148786 and ENSR00000251495 bivariate model of predicting irAEs. (R=0.79, P=$3.08\times10^{-6}$). The equation of the best bivariate model is 0.3732* ENSR00000148786+0.2181*ENSR-00000251495+1.2144. (**C**) Spearman correlation between ENSR00000187665 expression and ORR (R=0.59, *P*=7.5e-3). (**D**) Combined effect of ENSR00000035913 and ENSR00000167231 bivariate model of predicting ORR (R=0.82,

*P*=2.0e-5). The equation of the best bivariate model is 0.0063 * ENSR00000035913+0.1596 * ENSR00-000167231+0.1070.

**Fig. S1**

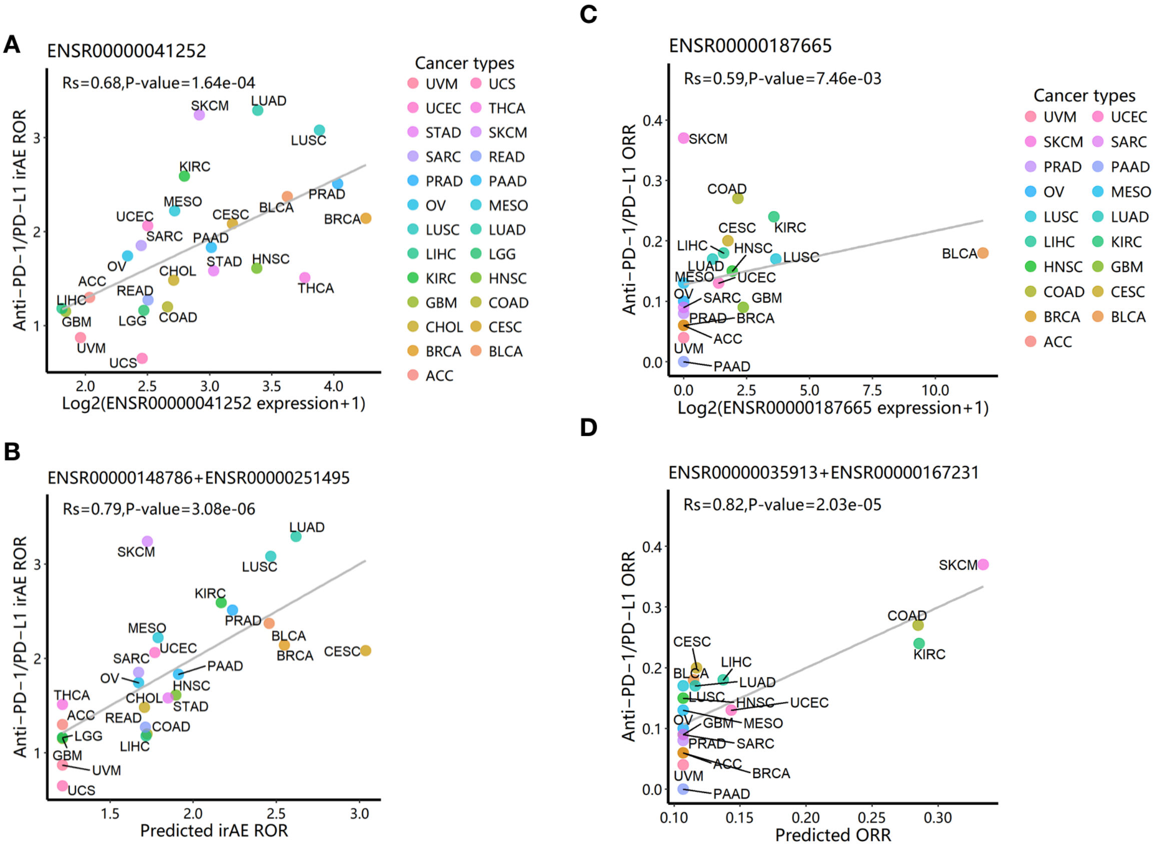

**Fig. S2. irAE-related genes involved in oncogenic pathways**. (**A**) RAF1 and RTK-RAS signaling pathway. (**B**) MLXIPL and MYC signaling pathway. Pathway graphs were generated by using the cBioPortal website.

#### Fig. S2

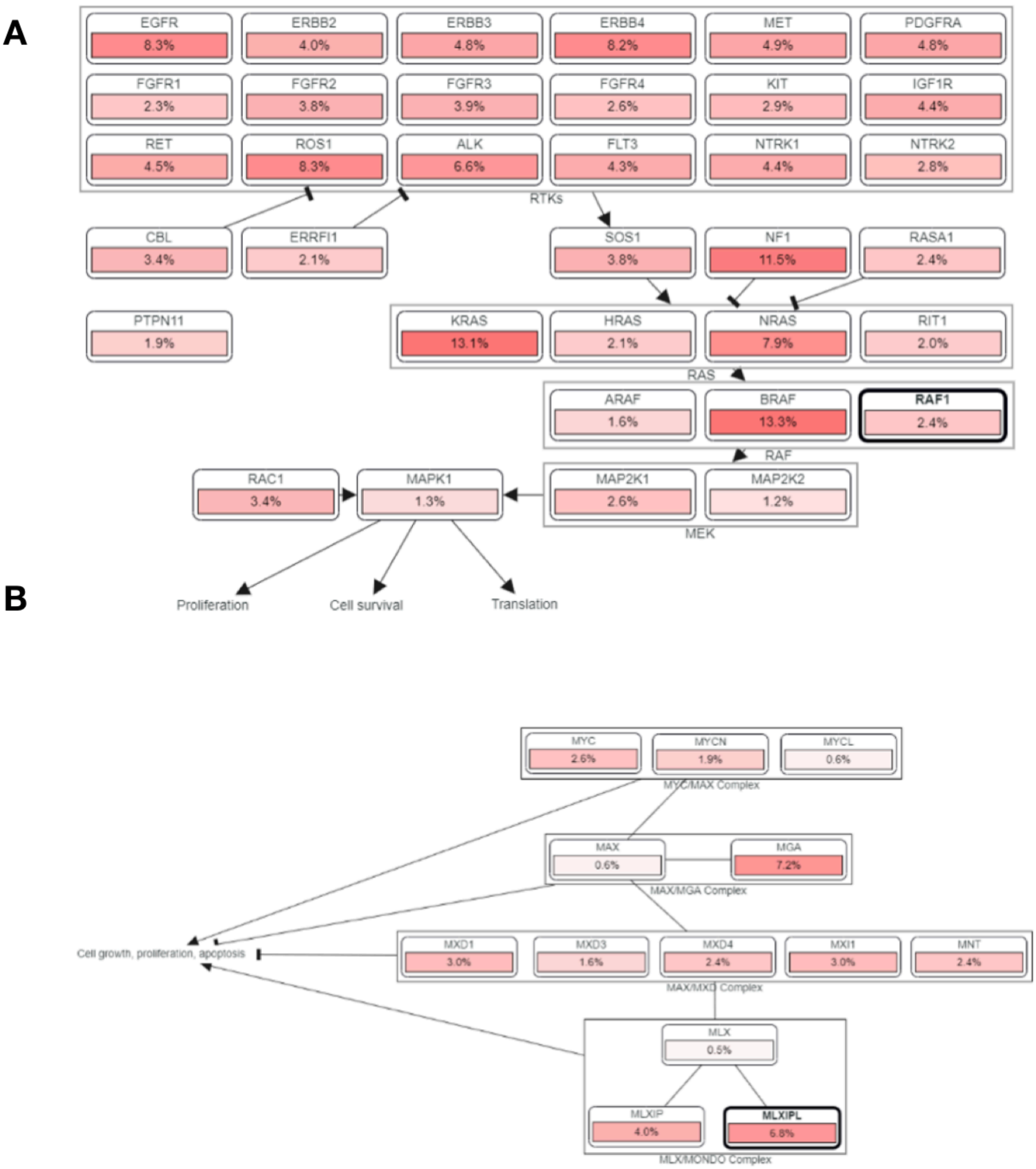
